# SHINE: Protein Language Model based Pathogenicity Prediction for Inframe Insertion and Deletion Variants

**DOI:** 10.1101/2022.08.30.505840

**Authors:** Xiao Fan, Hongbing Pan, Alan Tian, Wendy K. Chung, Yufeng Shen

## Abstract

Inframe insertion and deletion variants (indels) alter protein sequence and length. Accurate pathogenicity predictions are important in genetic studies of human diseases. Indel Interpretation is challenging due to limitations in the available number of known pathogenic variants for training. Existing methods largely use manually encoded features including conservation, protein structure and function, and allele frequency. Recent advances in deep learning modeling of protein sequences and structures provide an opportunity to improve the representation of salient features based on large numbers of protein sequences. We developed a new pathogenicity predictor for <u>SH</u>ort Inframe i<u>N</u>sertion and d<u>E</u>letion (SHINE). SHINE uses pre-trained protein language models to construct a latent representation of an indel and its protein context from protein sequences and multiple protein sequence alignments, and feeds the latent representation into supervised machine learning models for pathogenicity prediction. We curated training data from ClinVar and gnomAD, and created two test datasets from different sources. SHINE achieved better prediction performance than existing methods for both deletion and insertion variants in these two test datasets. Our work suggests that unsupervised protein language models can provide valuable information about proteins, and new methods based on these models can improve variant interpretation in genetic analyses.

## Introduction

Inframe insertion and deletion variants (indels) are abundant but are under studied in genetic analyses. In the ∼ 500,000 UK Biobank whole exome sequencing dataset, median numbers of the inframe and frameshift variants per individual were 115 and 90, respectively (1). A recent deep mutational scan study of *DDX3X* showed 49% of 625 inframe deletions caused cell depletion (2). While many inframe indels contribute to human diseases (3-5), the impact of rare individual inframe indels is usually uncertain. In ClinVar (6), 64% and 18% of ∼ 12,000 inframe indels are reported as variants of uncertain significance (VUS) and pathogenic/likely pathogenic (P/LP), compared with 12% and 86% of 53,000 frameshift indels are VUS and P/LP (7). This issue is due to the difficulty in determining pathogenicity of inframe indels, despite publications of several computational prediction methods (8-16).

Accurate pathogenicity prediction for inframe indels is limited by the availability of training data. There are only 2,208 P/LP and 1,599 benign/likely benign (B/LB) inframe indels with two stars (multiple submitters with assertion criteria and no conflicting interpretation) in ClinVar (6). Additionally, nearly all existing computational methods use machine learning with manually encoded predictive features, such as DNA/protein conservation, protein structure and function, occurrence in a repeat region, size of the indels, local amino acid composition, distance to the nearest splice site, and allele frequency. Although these features are correlated with variant pathogenicity, they are based on our limited understanding of how variants exert their function. Recent advances in deep learning, particularly protein language models, enable us to study millions of protein sequences through self-supervised learning (17-19). The models generate latent representations for protein sequences that can potentially capture salient information not encoded by existing methods. This opens up an opportunity to improve the input salient features used in protein-related bioinformatic applications.

Here we present SHINE (<u>SH</u>ort Inframe i<u>N</u>sertion and d<u>E</u>letion), a transfer learning model to leverage protein language models and limited available pathogenicity data for inframe indels. The protein language models take protein sequences or multiple protein sequence alignments (MSA), and generate latent representations capturing protein features. Previous studies have shown the linear projections of the representations generated from the protein language models encode information about protein secondary and tertiary structure (18). The same group used zero-shot and few-shot transfer of the protein language models to predict variant effects (20). SHINE uses the representations as features to separate pathogenic from benign inframe indels. It is trained on curated inframe indels from ClinVar, and short and common indels in gnomAD; and compared with the other computational methods on two independent test datasets. Finally, we interpreted the salient features from SHINE by comparing them with known predictive protein features, such as protein secondary structure, intrinsically disordered protein regions, and relative solvent accessibility.

## Materials and methods

### Data sets

We obtained P/LP and B/LB inframe indels from ClinVar (6) and gnomAD, and excluded inframe indels that have a length greater than three amino acids or allele frequency <0.1% in gnomAD. We divided this dataset into training and validation datasets based on the number of deleted/inserted amino acids. Indels with one amino acid deletion/insertion are used for training and the rest are for validation. The training dataset includes 1,040 pathogenic and 1,111 benign deletions, and 142 pathogenic and 537 benign insertions. The validation dataset includes 640 pathogenic and 896 benign deletions, and 272 pathogenic and 662 benign insertions. These datasets are used to select the optimal number of features, prediction models, and parameters for the models.

To evaluate the performance, we created two independent datasets for neurodevelopmental disorder (NDD) genes and cancer driver genes. Overlapping indels with the training-validation datasets were removed from the test datasets. The two test datasets are from different sources than ClinVar, which avoids inflated performance when testing on data from the same source. Besides, the two datasets were not used to train any of the previous prediction models for indels, so are good benchmarks for method comparison. Six hundred and eighty-six NDD risk genes were collected based on previous studies (21,22). *De novo* inframe indels in the 686 NDD genes in two NDD cohorts were used as proxies of pathogenic variants, and short inframe indels with at most three-amino-acid deletion/insertion in the UK biobank (200,000 exomes) were considered benign. One NDD cohort includes 16,877 trios in Simons Powering Autism Research (SPARK) (21,23,24) and the other includes 13,058 trios with developmental disorders (22). This test dataset includes 146 pathogenic and 2,808 benign deletions, and 35 pathogenic and 1,504 benign insertions. The second test dataset is composed of inframe indels discovered in more than 1,000 cancer mutational hotspots (25). We identified 307 deletions and 119 insertions in 36 genes as proxies of pathogenic variants. We extracted short inframe indels including 132 deletions and 54 insertions from the UK biobank in the same 36 genes as benign. To be noted, the variants from both cases and controls are mixtures of pathogenic and benign variants, but the *de novo*/somatic variants in NDD risk genes/cancer mutational hotspots identified in cases are much more likely to be pathogenic compared with germline variants in controls. We used the case/control status as a proxy of pathogenic/benign labels in the test.

### Model architecture

SHINE uses a transfer learning architecture (Figure 1), leveraging pretrained protein language models and limited available pathogenicity labels for inframe indels. We used two protein language models: ESM-1b (18) and MSA transformers (27). The ESM-1b transformer was trained on 250 million protein sequences (18). The MSA transformer was trained on 26 million MSAs (27). Both generated latent representations containing information about biological properties of input proteins. We elaborate on how we selected proper transformers, ways to handle multiple-amino-acid indels, input dimensions (number of features), prediction models and their parameters. As the latent representations from the transformers are high dimensional and correlated, we first performed feature reduction using principal components analysis (PCA) on the 1,024 and 768 latent representations from the ESM-1b and the MSA transformer. The optimal number of remaining principal components (nPCs) was selected using linear regression as a base predictor. The transformed PCs were fed as salient features to a supervised machine learning model. We further tested different supervised machine learning models including random forest, support vector machine, gradient boosting and elastic net. The pre-selected optimal nPCs were used. Their optimal parameters were selected based on 5-fold cross validation on the training dataset using regression coefficients as optimized scores. Finally, we repeated the selection of the nPCs for each model given the tuned parameters. The above process was done on our training-validation datasets using sklearn (26) for deletions and insertions separately. For multiple-amino-acid indels, we calculated the predictive scores for each amino acid and then tested the maximum, mean and sum of the scores as the final predictive score.

**Figure 1.**
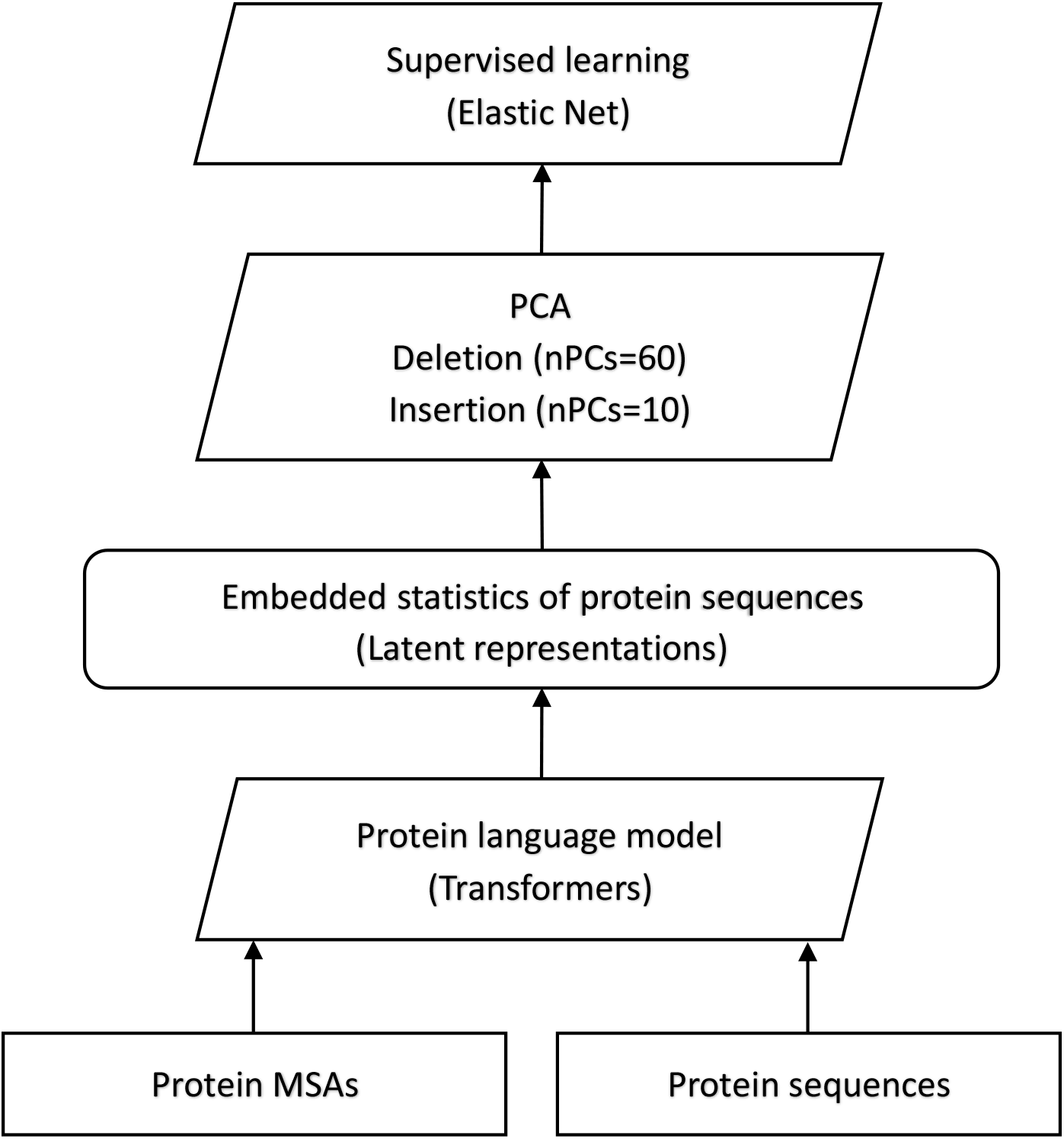
Architecture of the SHINE.

### Input features

ESM-1b and MSA transformers take protein primary sequences and MSAs as inputs. We downloaded MSA data from Ensembl Compara using REST API (https://rest.ensembl.org/documentation/info/genetree). The median and mean MSA depth is 211 and 320.2, respectively. As many proteins included in the MSA are not similar enough to the human proteins of interest, we trimmed the phylogenetic trees to remove the less similar proteins and included a maximum of 300 proteins per MSA. This also speeds up the pretraining process to generate latent representations. Supplementary Figure 1 shows the distribution of MSA depth before and after the trim. The median and mean MSA depth after the trim is 199 and 184.3, respectively. We fed wild-type protein sequences or MSAs to the pretrained transformers, and extracted latent representations of amino acids that were deleted. For insertions, wild-type MSAs were used for the MSA transformer. Extracted were latent representations of the position (amino acid or gap) followed by the amino acid where the insertion occurs. Mutated protein sequences with inserted amino acid(s) were inputted for ESM-1b transformer and latent representations of the inserted amino acid(s) were used as features. The dimensions of the latent representations from the two transformers are 1024 and 768, respectively.

### Measures of predictive quality

Quantitative predictive scores were evaluated using the Receiver Operating Characteristic (ROC) curve and the difference of two score distributions from cases and controls. R package ‘pROC’ was used to estimate Area Under the ROC Curve (AUC) and to evaluate the level of significant difference between two ROC curves. AUC is used as the quality measure to select the optimal number of features, models, and way to handle multiple-amino-acid indels. The difference between the two distributions from cases and controls was measured using Wilcoxon rank sum test in R. Smaller *P*-values indicate better separation of benign and pathogenic variants. The binary predictions were assessed using balanced accuracy, sensitivity/recall, and specificity.

### Existing pathogenicity predictors for inframe indels

We selected all available methods that predict pathogenicity for inframe indels and provide a web server for convenient access to the end users. They included VEST-indel (12), CAPICE (16), and CADD (15). VEST-indel applied random forest included 24 features. CADD used L2 regularized logistic regression with more than 60 features. CAPICE used gradient boosting on decision trees with the same set of features as CADD. The default thresholds for SHINE, CAPICE, VEST and CADD are 0, 0.02, 0.8 and 20, respectively. Indels with predictive scores larger than the threshold are considered as pathogenic.

## Results

### SHINE model

Using transfer learning, SHINE takes advantage of the pretrained protein language models to maximize the use of limited available pathogenicity labels for inframe indels. We first tested the performance of the two pretrained models: ESM-1b and MSA transformers. PCA was applied to their latent representations separately and then to combined representations to generate different sizes of salient features (nPCs). Linear regression projecting the salient features to pathogenicity label, was trained on the training dataset and tested on the validation dataset. We picked the highest (worst) predictive score for multiple-amino-acid insertions/deletions as the final predictive score. Supplementary Figure S2 shows the AUC values for the MSA transformer are higher than those for the ESM-1b transformer. Combining the representations from both models does not improve predictions compared with the MSA transformer only. The two transformers capture different properties about proteins: representations from the ESM-1b transformer consider amino acid sequence patterns and representations from the MSA transformer consider cross-species conservation. Therefore, we used both transformers in SHINE.

The following analyses were performed using representations from both ESM-1b and MSA transformers. We evaluated different ways (maximum, mean and sum) to handle multiple-amino-acid indels. Supplementary Figure S3 shows maximum score is clearly better. Although intuitively, long indels are more likely to be pathogenic than short indels, sum of the individual scores for each amino acid does not work. Thus, SHINE model uses maximum score for multiple-amino-acid indels.

PCA transformed the combined latent representations from the two transformers into low dimensional features (PCs). Figure 2 shows the scatterplot of the first two PCs for pathogenic and benign variants. PC1 and PC2 correlated with pathogenicity for deletions with correlation coefficients of -0.376 and -0.464. The correlation coefficients for insertions were 0.482 and - 0.415 for PC1 and PC2, respectively. The first 10 PCs explained 41.0% and 40.3% of the variance for deletion and insertion representations, respectively. Scree plots are in Supplementary Figure S4. Eighty and ten PCs were chosen for deletions and insertions as they give the highest AUC values based on the linear regression model (Supplementary Figure S3).

**Figure 2.**
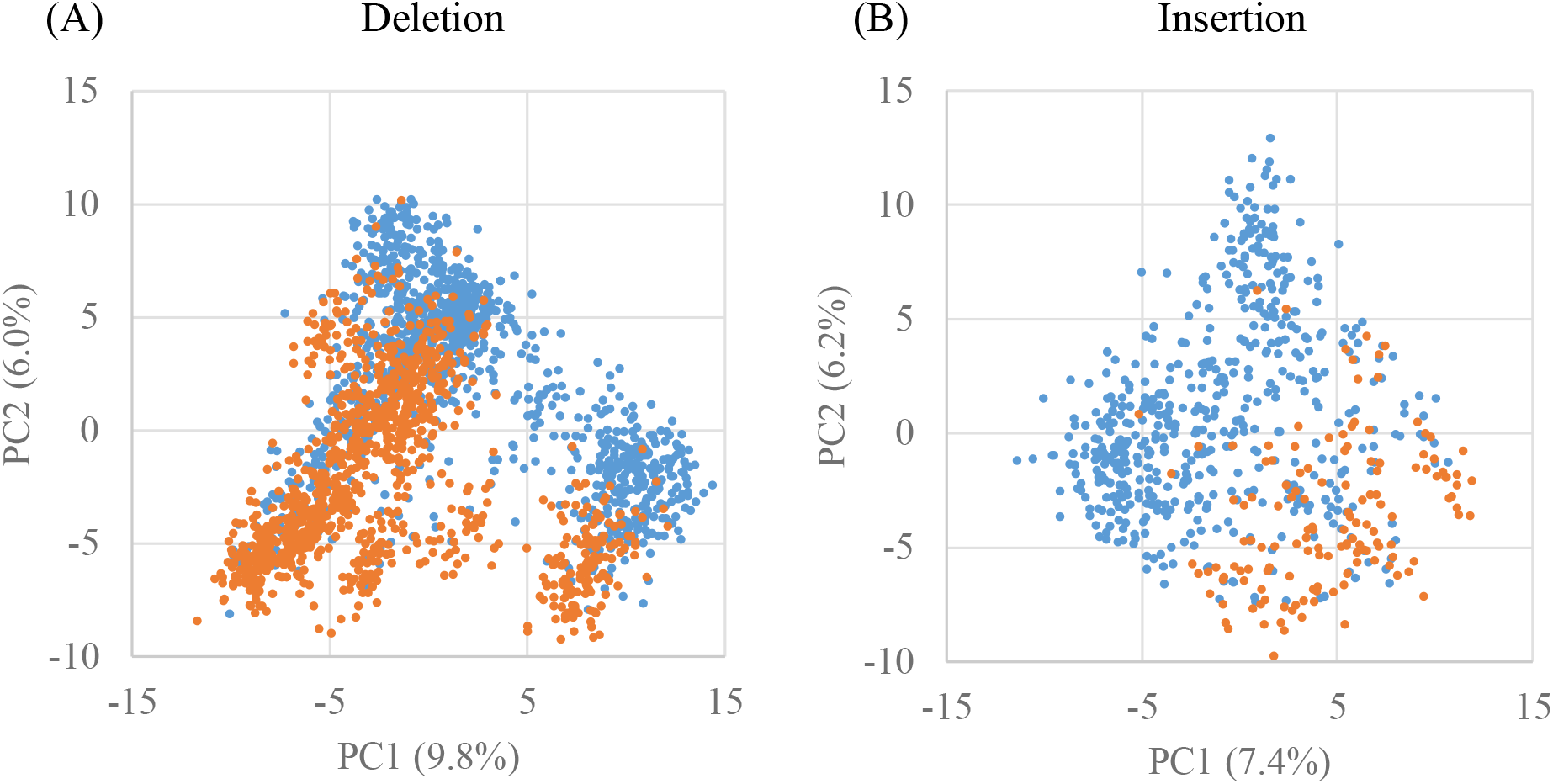
Scatterplot of the first two principal components for pathogenic and benign inframe indels.

We tested different supervised machine learning models and tuned their parameters given the same nPCs optimized in the last step as input. At the end, each of the tuned models was used to select their own optimal nPCs based on the best AUC values. Elastic Nets were the best models as they provided consistent good performance, were not sensitive to the number of input PCs and were not likely to overfit on the training dataset (Figure 3). Parameters alpha and l1_ratio of the Elastic Nets were 0.5 and 0.1 for both deletions and insertions. The Elastic Net for the deletions included 60 PCs and 24 of them had non-zero coefficients. The Elastic Net for the insertions included 10 PCs, and all of them had non-zero coefficients.

**Figure 3.**
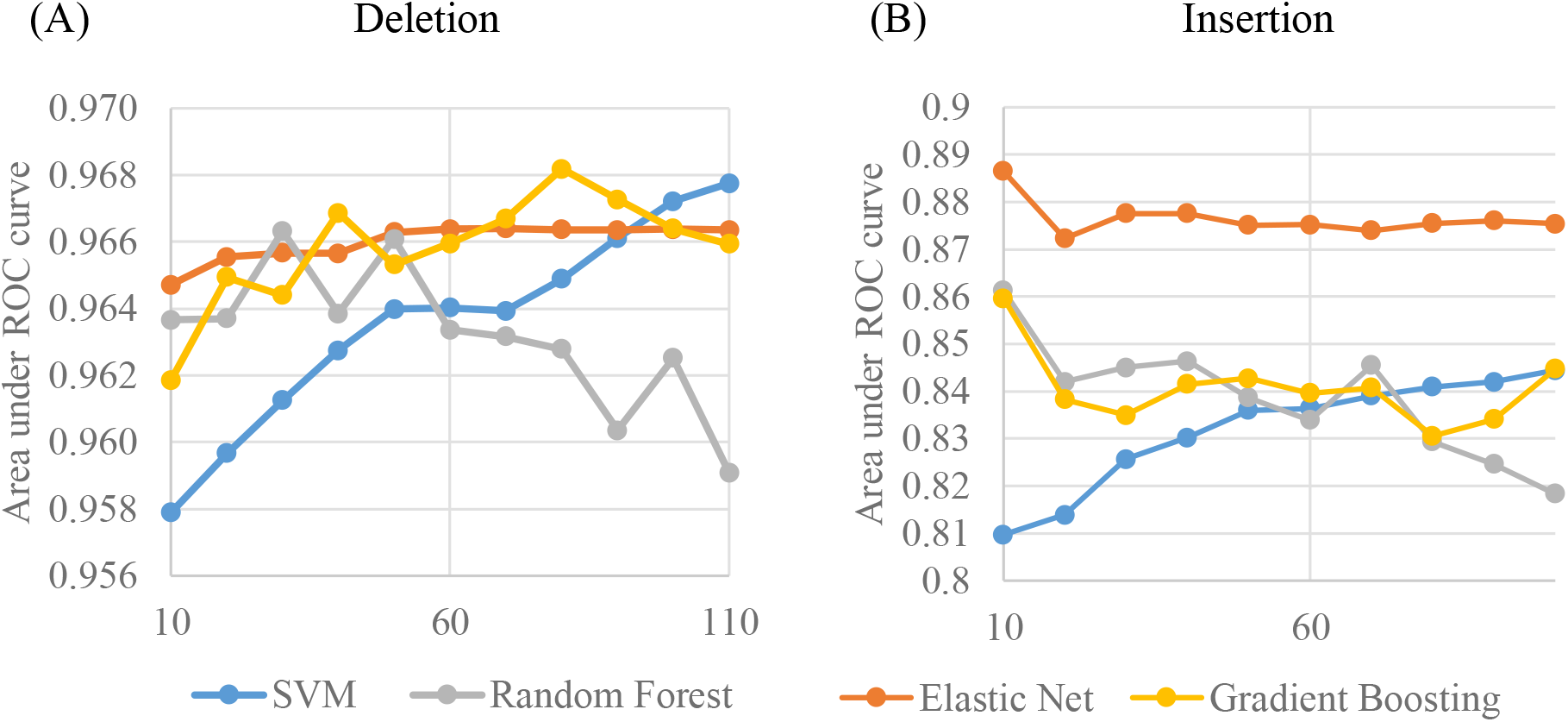
Area under ROC curve for a set of machine learning models with varying numbers of principal components as input features.

### Evaluation on the NDD test dataset

Instead of splitting data for training and test from the same source, we used independent test datasets to avoid the inflated performance on data from the same source. We compared SHINE with three methods including VEST-indel (12), CAPICE (16) and CADD (15) by classifying indels in the 686 NDD risk genes from NDD cases and UK biobank controls. We note the indels from both cases and controls are mixtures of pathogenic and benign variants, but the *de novo* variants in NDD risk genes identified in cases are more likely pathogenic compared with germline variants in the general population. Thus, we could use the case/control status as a proxy of pathogenic/benign labels in the test. We first examined the predictive power of the autism (SPARK) and NDD cohorts separately. Supplementary Figure S5 shows SHINE scores have distinct modes for cases versus controls for both cohorts, so we combine autism and NDD cases for the following analysis. Compared with the other computational methods, SHINE has the lowest *P*-value (Supplementary Figure S6), i.e., separates pathogenic from benign the best.

From the ROC curves, Figure 4 shows that SHINE has the highest AUC values of 0.846 and 0.834, improves over the second-best VEST-indel by 0.04 (relative improvement of 4.4%) and 0.07 (relative improvement of 8.5%) for deletions and insertions, respectively. The improvement over all other three methods is significant (*p*-value<0.05). The sensitivity rises quickly at a low false positive rate for SHINE. It implies SHINE scores correlate with likelihood of pathogenicity. We evaluated their binary predictions using their default thresholds. Supplementary Table S2 shows the evaluation for the binary prediction including balanced accuracy, sensitivity and specificity. SHINE provides the best balanced accuracy scores of 0.769 and 0.748 for deletions and insertions. All the methods over-predict. CAPICE predicts the most, so has the highest sensitivity. SHINE, VEST and CADD offer the best specificity. Overall, SHINE and VEST give good balanced accuracy balancing sensitivity and specificity. Our analysis suggests SHINE can distinguish pathogenic from benign indels well. For users interested in high-precision predictions, high SHINE scores provide a good solution (high sensitivity given a low false positive rate in Figure 4).

**Figure 4.**
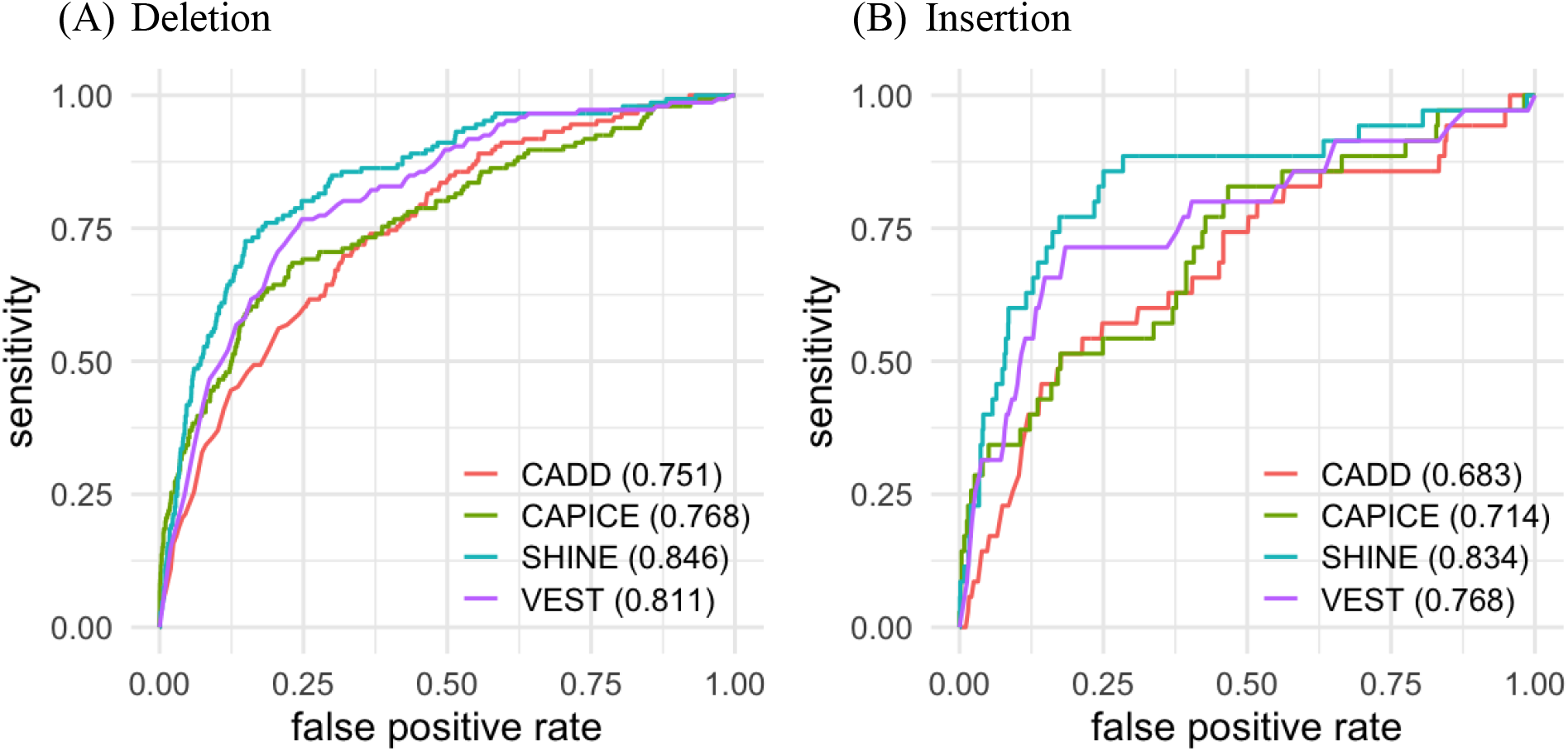
Evaluation of predictive scores using receiver operating characteristic curves for four prediction methods in NDD cases and UK biobank controls.

None of the computational methods used allele frequency in their predictive models, so we tested their sensitivity to allele frequency. We repeated the analysis using a subset of benign variants with gnomAD allele frequency > 10^−4^ (146 deletions and 35 insertions in cases, and 601 deletions and 339 insertions in controls). All the computational methods perform better on this dataset with more restricted benign indels. SHINE still has the highest AUC value (Supplementary Table S1).

One-amino-acid inframe indels are most abundant among indels in the population and are also the hardest to predict. We extracted indels from our NDD test dataset with only one deleted or inserted amino acid. This subset includes 118 deletions and 18 insertions in cases, and 2,207 deletions and 747 insertions in controls. We reported the AUC values on this subset in Supplementary Table S1. All methods provide similar AUC values compared with that from the full NDD dataset.

### Evaluation on the cancer mutational hotspot test dataset

We compared SHINE with the other three methods on indels in the other independent test dataset: cancer mutational hotspots and UK biobank. Supplementary Figure S7 shows SHINE has the lowest *P*-value to separate distributions of predictive scores for cases and controls. VEST provides the same performance for insertions, but its performance on deletions is inferior. AUC results are consistent (Figure 5). SHINE with an AUC value of 0.882, is significantly better than VEST and CADD for deletions (*P*-value < 0.05). It improves over CAPICE by 0.04 (relatively improvement of 4.3%), but the difference is not significant. On the insertion dataset, SHINE and VEST with AUC values of 0.932 are insignificantly different. Both outperform the other two methods significantly by 0.17 (22.2%) in AUC values. All in all, our method is consistently the best on the two independent test datasets, despite different disease types.

**Figure 5.**
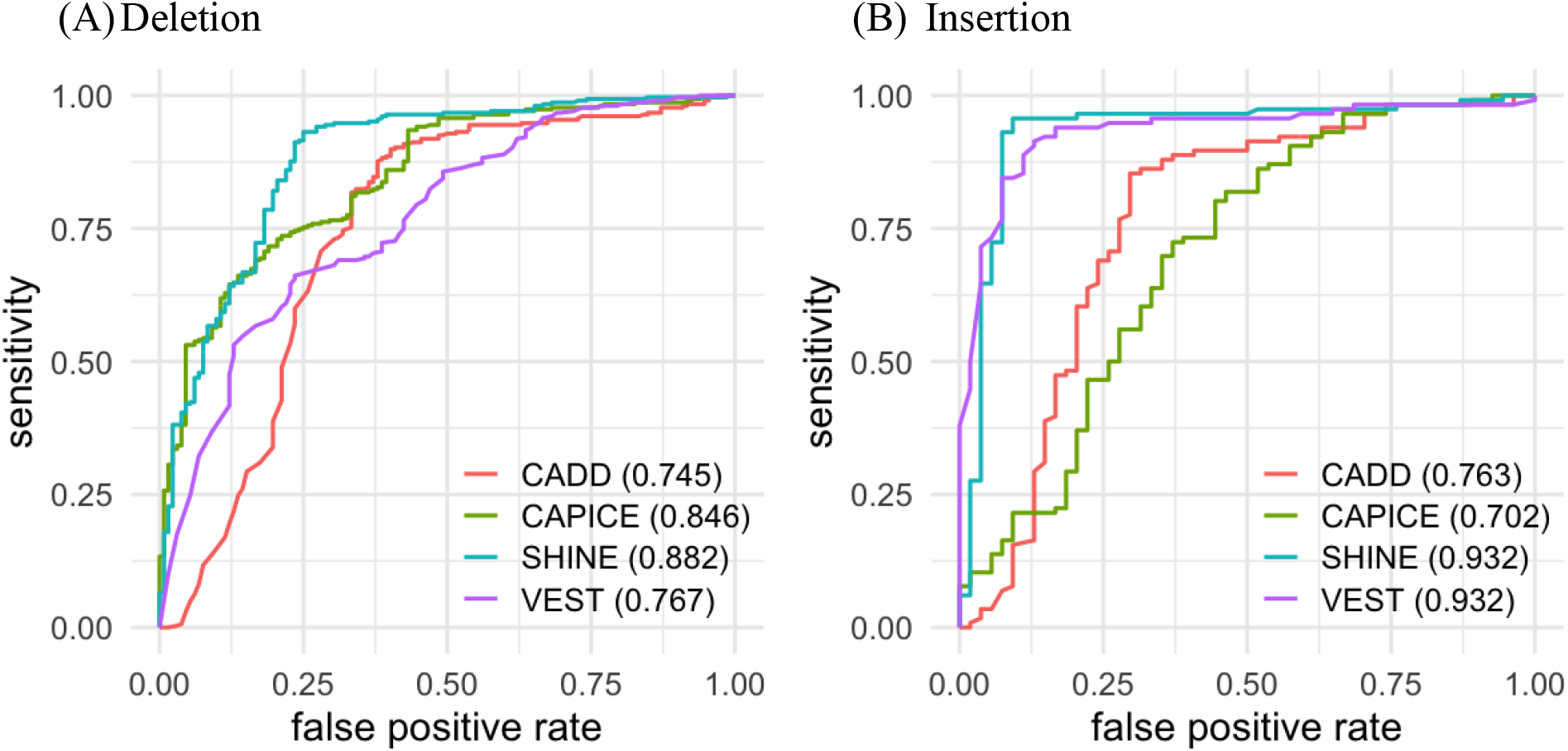
Evaluation of predictive scores using receiver operating characteristic curves for four prediction methods in cancer mutational hotspot cases and UK biobank controls.

### Interpretation of the input features

Previous studies identified a few top performing features for indel pathogenicity prediction, such as intrinsically disordered protein region, protein secondary structure and solvent accessible surface area (9). We investigated the correlation between the input features of SHINE – principal components and the three protein-structure features. Supplementary Figure S8 shows the PC1 is significantly correlated with protein secondary structure (coil), intrinsically disordered residues, and relative solvent accessibility. Pathogenic variants are more likely to be seen in structured protein regions and embedded in the core of proteins. Our observations are consistent with the previous studies (9). Even though we did not encode protein features as previous methods, these features are likely captured in the latent representations generated from the pretrained transformers. Moreover, the latent representations have potential to carry information that we have not discovered. More efforts are still needed to interpret the protein structure and function to better understand how genetic variations affect the proteins.

## Discussion

Accurate pathogenicity predictions for inframe indels are important. Available computational approaches to address this question are insufficient, primarily due to the limitations of available pathogenicity labels for training. We take advantage of the protein language models trained using millions of protein sequences to learn protein statistics/features in an unsupervised fashion. We for the first time transfer the knowledge about proteins to pathogenicity prediction for inframe indels. Evaluating on two independent test datasets, our method is consistently better than the current computational methods. SHINE scores provide a pathogenicity likelihood which can facilitate selection of top pathogenic inframe indels in research and clinical interpretation of variant pathogenicity.

Most of the current computational studies use pathogenic indels from ClinVar and Human Gene Mutation Database, and common and low-frequency indels from population datasets as benign. The issue is that pathogenic indels have been identified only in a few hundred known disease genes, while benign indels are largely extracted from unconstrained genes. The two sets of genes do not overlap much. Our training-validation datasets include pathogenic indels in 917 genes and benign indels in 1,566 genes, and only 206 genes contributing both pathogenic and benign indels. Previous methods built on similar datasets tended to select gene-level features which separate disease genes from the rest. These methods lack the power to distinguish pathogenic from benign indels when testing the same gene. Our test datasets are designed to eliminate the effect from the ascertainment bias. We included variants in the same set of genes from different sources. Neither the NDD nor the cancer mutational hotspot dataset was used to train any of the previous methods. So, our test datasets are good benchmarks to test the predictive power for variant interpretation and compare across different approaches.

We note the labels in our test datasets are not true pathogenic or benign classifications, instead case/control status for the origin of the variants. Not all variants found in cases are pathogenic; not all variants found in controls are benign. In our sensitivity analysis removing ultra-rare variants from controls, all methods perform better. It suggests it is easier to separate pathogenic variants from common benign variants compared with rare benign variants. We also acknowledge a small portion of rare variants in controls could be pathogenic, which affects the predictive performance of the computational methods.

The protein representations learnt from the pretrained protein language models correlate with the previously discovered predictive features and provide better predictive power. Moreover, these representations are protein signatures and not biased to any specific task, so can be used for many other bioinformatic applications.

## Conclusions

SHINE is the first protein language model-based method to predict pathogenicity of inframe indels. The protein language models generate unbiased protein statistics in an unsupervised fashion. Future research should consider expanding the variant types for pathogenicity prediction using similar approaches. Benchmark datasets from functional data will be highly appreciated as deep mutational scan data are becoming available for inframe indels.

## Data availability

The training, validation and test datasets, and codes for processing the MSA data and running the prediction models are available on GitHub: https://github.com/xf-omics/SHINE

## Acknowledgement

We thank Dr. Chen Wang for helpful discussions. We are grateful to all of the families in SPARK study, the SPARK clinical sites and SPARK staff. We appreciate obtaining access to the genomic data on SFARI Base. Approved researchers can obtain the SPARK population dataset described in this study by applying at https://base.sfari.org. We appreciate obtaining access to recruit participants through SPARK research match on SFARI Base. The SPARK initiative is funded by the Simons Foundation as part of SFARI.

## Author contributions

Conceptualization: X.F., Y.S.; data curation: X.F., H.P.; formal analysis and visualization: X.F.; writing manuscript: X.F., Y.S., W.K.C.; supervision: Y.S., W.K.C.; All authors reviewed the manuscript and approved the submission of this manuscript.

