## Supplemental Data for "SHINE: Protein Language Model based Pathogenicity Prediction for Inframe Insertion and Deletion Variants"

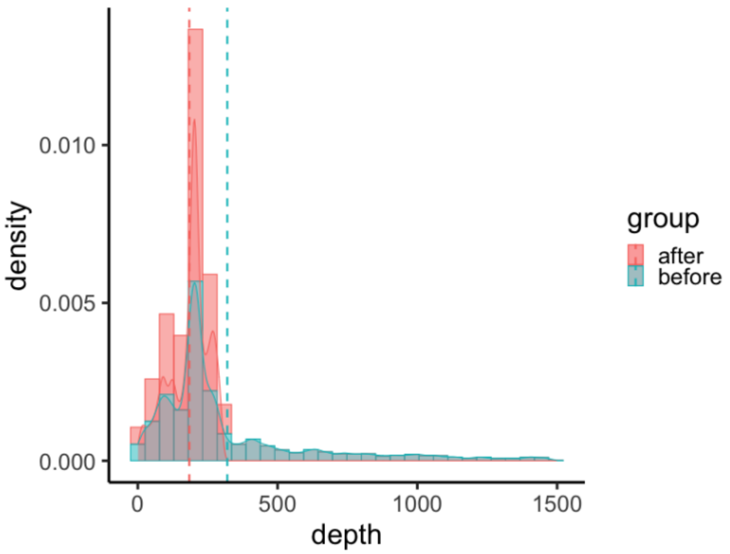


**Figure S1.** Density of multiple sequence alignment depth before and after trim of phylogenetic tree.

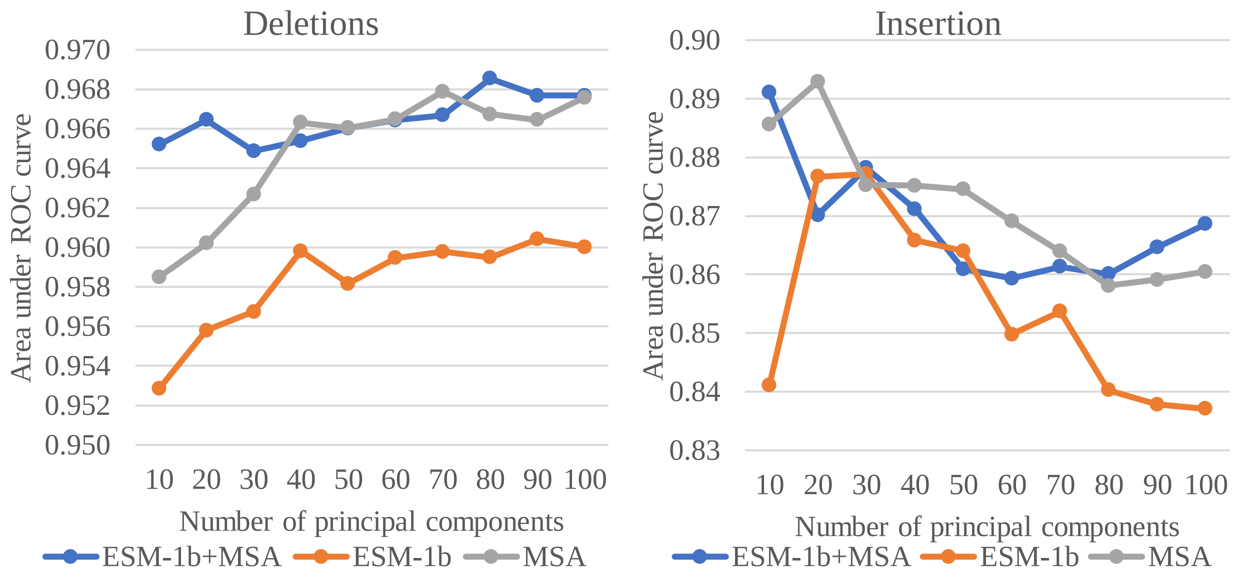


**Figure S2.** Area under the ROC curve with varying numbers of principal components as input features for different transformer models or combination.

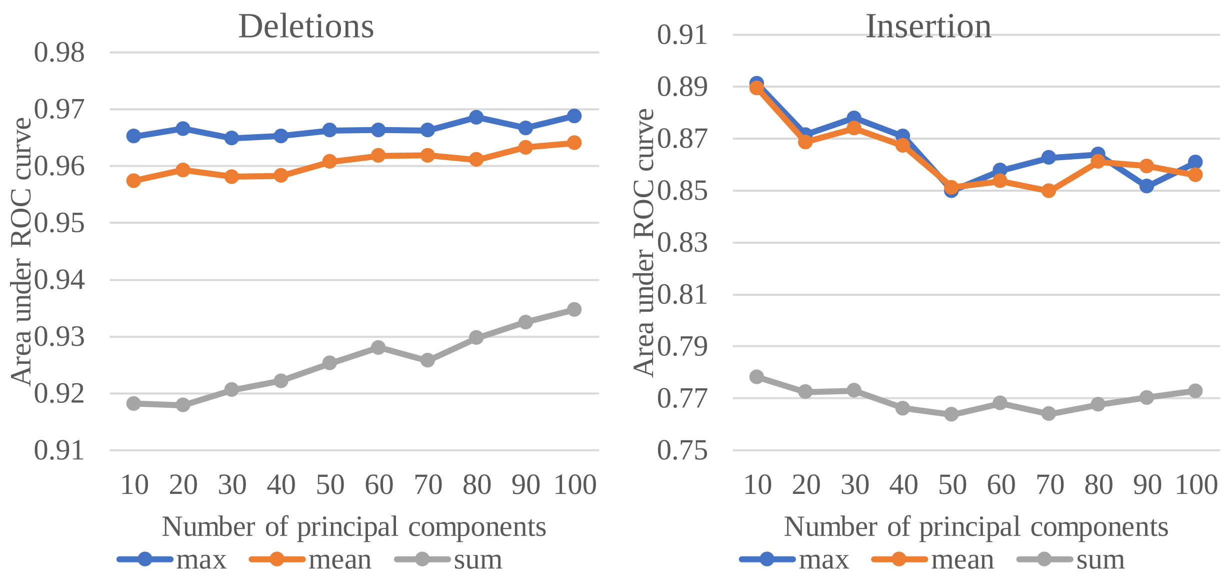


**Figure S3.** Area under the ROC curve using different ways to handle the multiple amino acid indels.

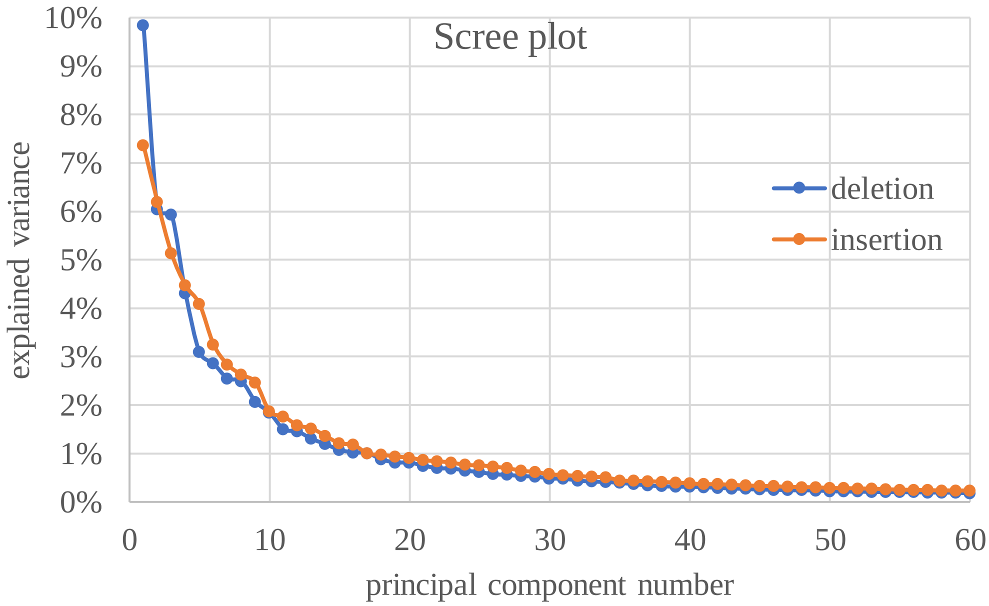


**Figure S4.** Scree plot for variance explained by individual principal components on the training dataset.

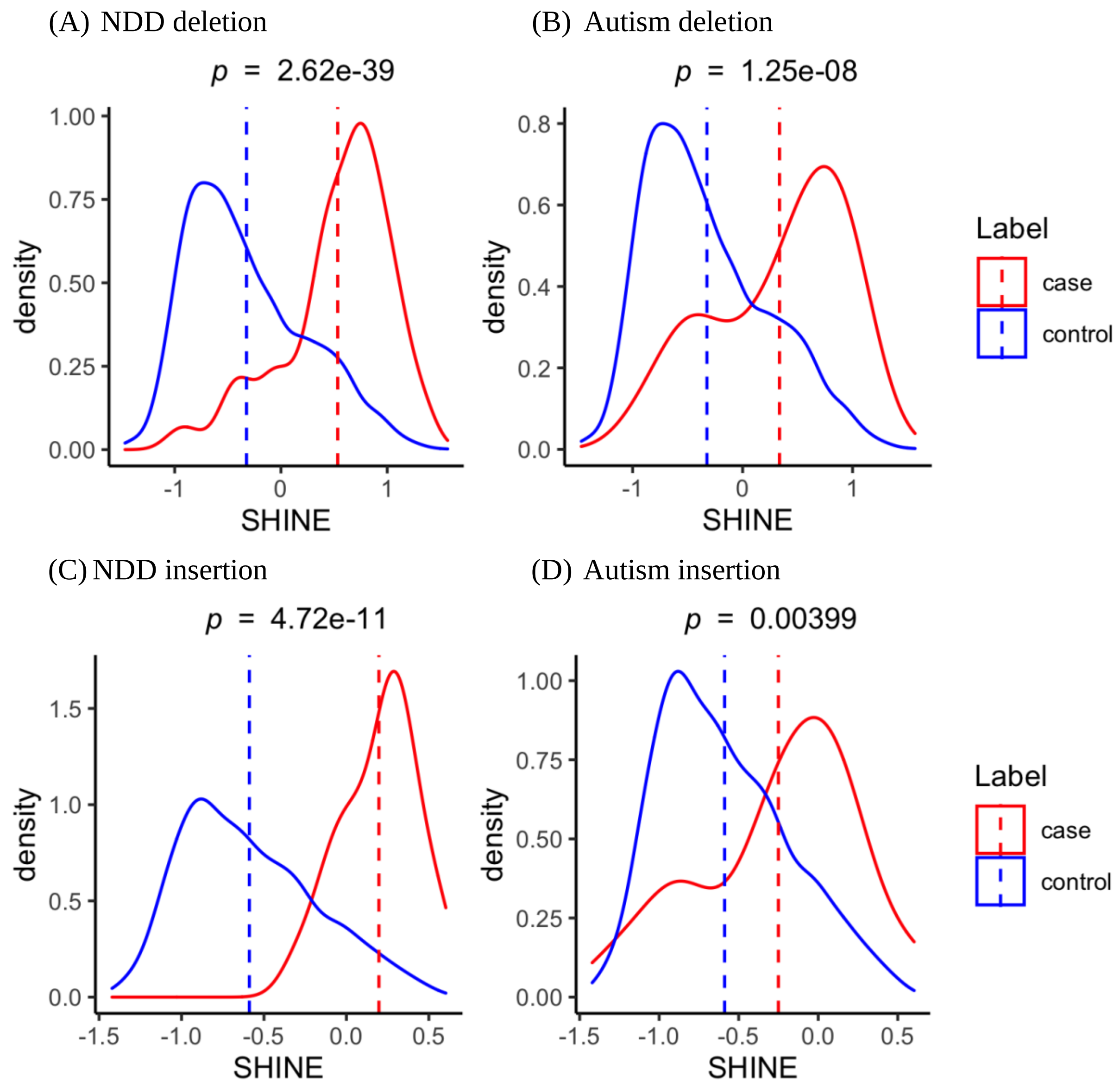


**Figure S5.** Distributions of SHINE scores of inframe deletions/insertions in NDD and Autism cohorts.

(A) Deletion (B) Insertion

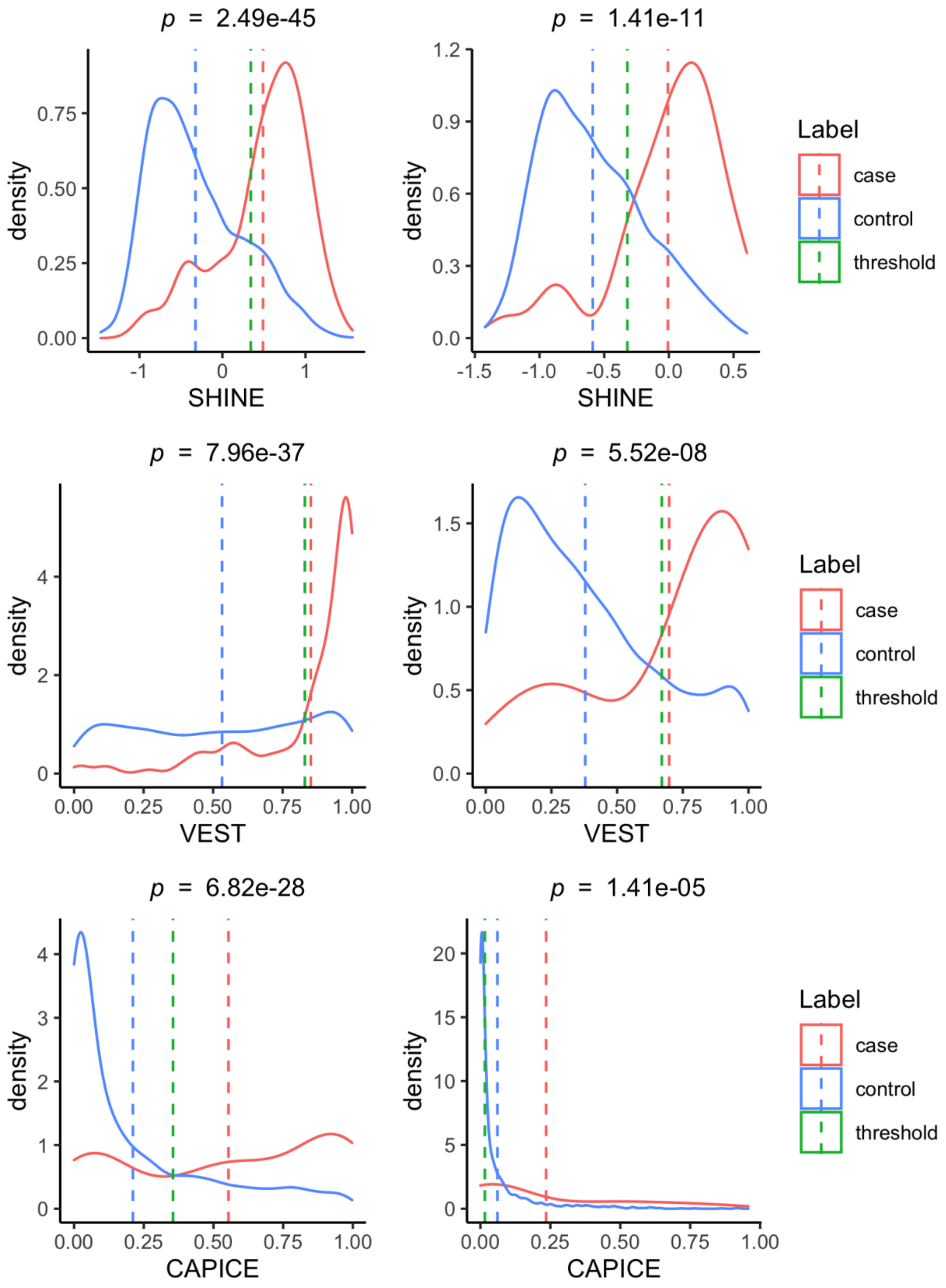

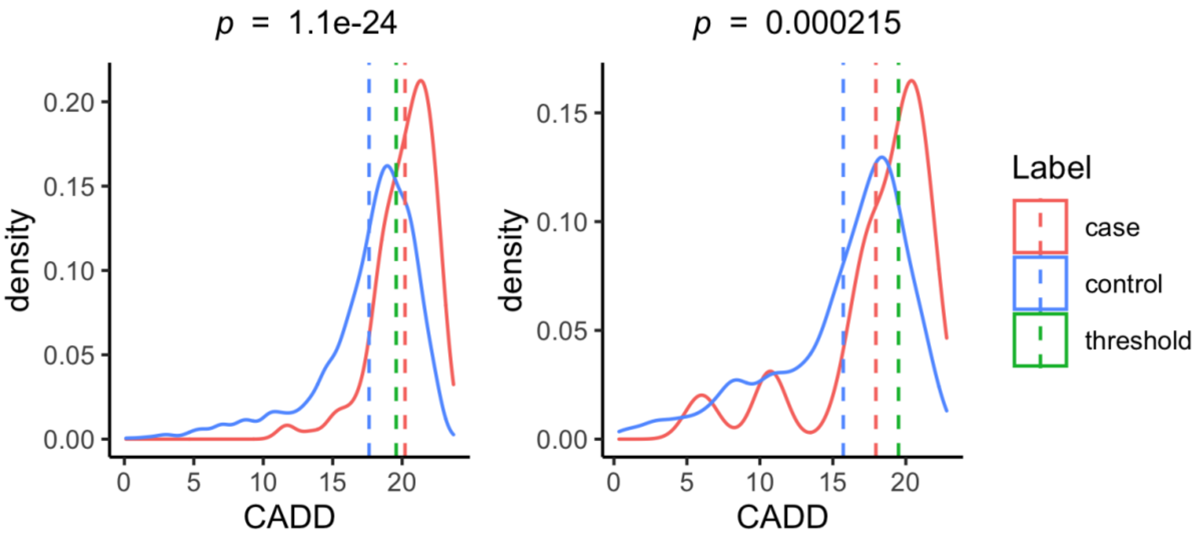


**Figure S6.** Distribution of predictive scores in NDD cases (red) and UK biobank controls (blue). Red and blue dashed lines indicate the mean predictive scores for cases and controls, respectively. Green dashed line marks the optimal threshold of predictive scores to separate cases from controls given the best balanced accuracy value.

(A) Deletion (B) Insertion

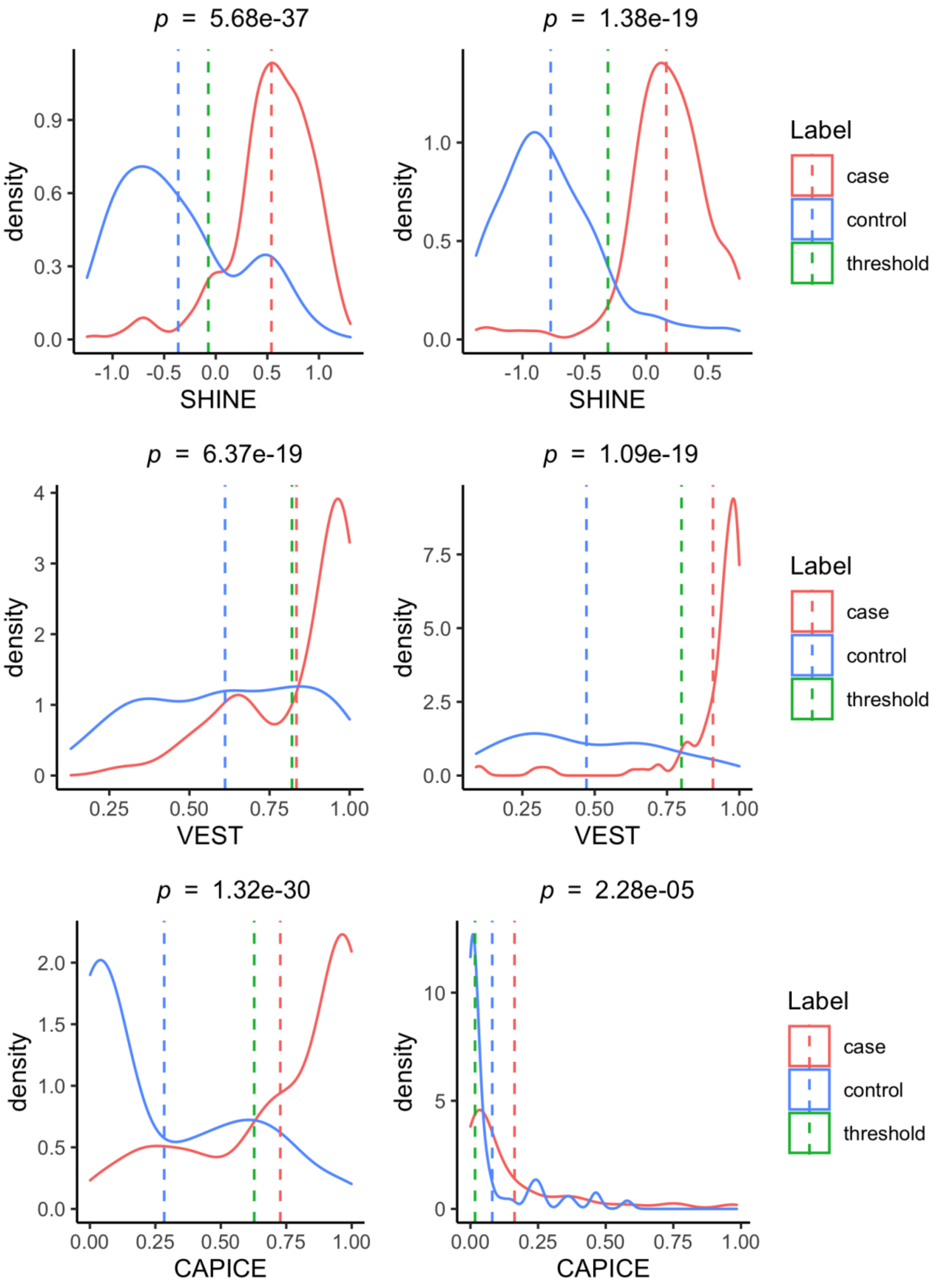

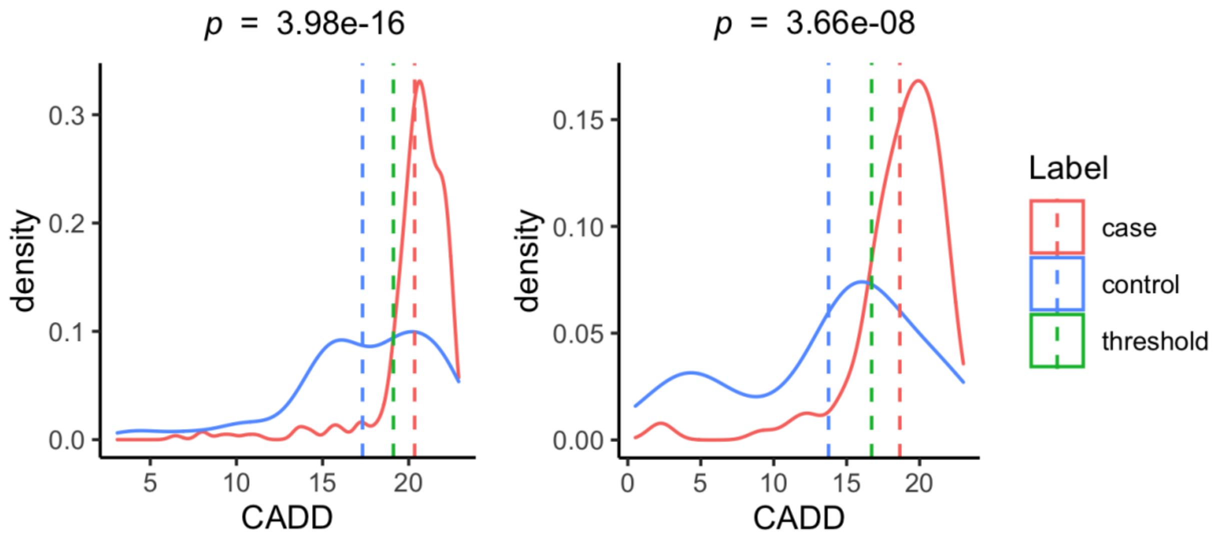


**Figure S7.** Distribution of predictive scores in cancer mutational hotspots (red) and UK biobank controls (blue). Red and blue dashed lines indicate the mean predictive scores for cases and controls, respectively. Green dashed line marks the optimal threshold of predictive scores to separate cases from controls given the best balanced accuracy value.

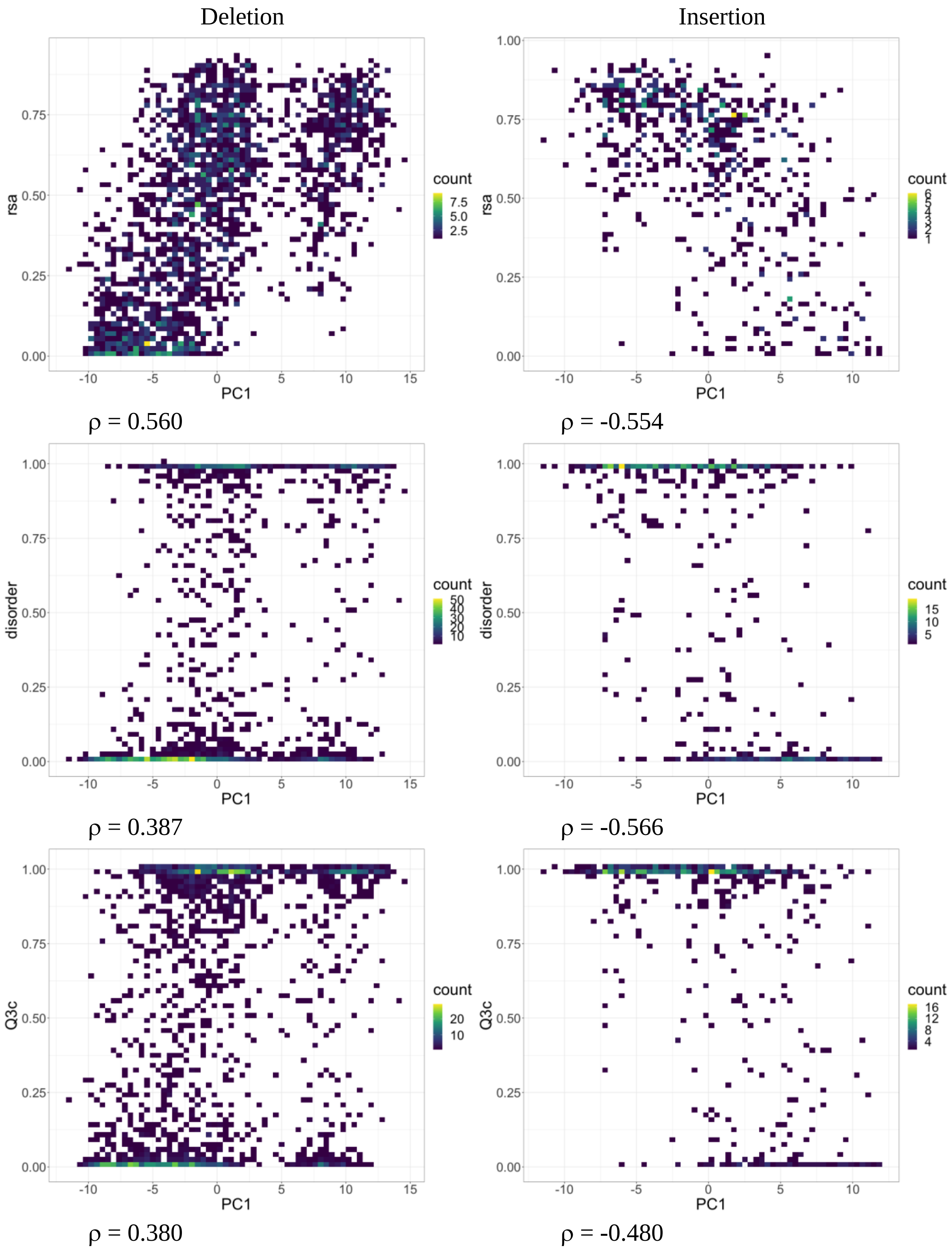
Deletion
Insertion
rsa
PC1
count
ρ = 0.560
ρ = -0.554
disorder
PC1
count
ρ = 0.387
ρ = -0.566
Q3c
PC1
count
ρ = 0.380
ρ = -0.480

**Figure S8.** Distribution of the first principal component (PC1) values and protein properties, from the top: relative solvent accessibility (rsa), intrinsically disordered residues (disorder), and coil in protein secondary structure (Q3c). The correlation coefficients are shown below each plot.

**Table S1.** Area under ROC curve for inframe indels in NDD cases and UK biobank controls. Numbers of pathogenic (P) and benign (B) variants are given in parentheses for each dataset. *Methods that are insignificantly different compared with SHINE.

| | **All** | | **Benign gnomAD AF>$10^{-4}$** | | **One amino acid** | |
|---|---|---|---|---|---|---|
| | Deletion (146P+2808) | Insertion (35P+1504B) | Deletion (146P+601B) | Insertion (35P+339) | Deletion (118P+2207B) | Insertion (18P+747B) |
| **SHINE** | **0.846** | **0.834** | **0.936** | **0.883** | **0.838** | **0.822** |
| **VEST** | 0.811 | 0.768 | 0.911 | 0.821 | 0.811* | 0.765* |
| **CADD** | 0.751 | 0.683 | 0.829 | 0.736 | 0.744 | 0.601 |
| **CAPICE** | 0.768 | 0.714 | 0.838 | 0.765 | 0.755 | 0.721* |

**Table S2.** Evaluation of binary predictions from four different methods. The default thresholds are used to generate the binary predictions.

| | Balanced accuracy | Sensitivity | Specificity | Threshold |
|---|---|---|---|---|
| **Deletion** | | | | |
| SHINE | **0.769** | 0.801 | **0.736** | 0 |
| VEST | 0.751 | 0.767 | 0.735 | 0.8 |
| CADD | 0.675 | 0.603 | **0.748** | 20 |
| CAPICE | 0.591 | **0.911** | 0.272 | 0.02 |
| **Insertion** | | | | |
| SHINE | **0.748** | 0.600 | **0.897** | 0 |
| VEST | 0.715 | 0.543 | 0.888 | 0.8 |
| CADD | 0.635 | 0.400 | 0.871 | 20 |
| CAPICE | 0.659 | **0.743** | 0.574 | 0.02 |
